# Toxigenic Profile of *Clostridium perfringens* Strains Isolated from Natural Ingredient Laboratory Animal Diets

**DOI:** 10.1101/2021.03.03.433836

**Authors:** Michael D. Johnston, Tanya E. Whiteside, Michelle E. Williamson, David M. Kurtz

## Abstract

*Clostridium perfringens* is an anaerobic, gram-positive, spore-forming bacterium that ubiquitously inhabits a wide variety of natural environments including the gastrointestinal tract of humans and animals. *C. perfringens* is an opportunistic enteropathogen capable of producing at least 20 different toxins in various combinations. Strains of *C. perfringens* are currently categorized into seven toxinotypes (A, B, C, D, E, F & G) based on the presence/absence of four major toxins (alpha, beta, epsilon & iota) and two minor toxins (enterotoxin & netB). Each toxinotype is associated with specific histotoxic and enteric diseases. The Quality Assurance Laboratory (QAL) at the National Institute of Environmental Health Sciences (NIEHS) screens incoming animal feeds for aerobic, enteric pathogens, such as *Salmonella* spp. and *E. coli*. Recently, QAL has incorporated anaerobic screening of incoming animal feeds. To date, the lab has isolated numerous *Clostridiu*m species, including *C. perfringens*, from 23 lots of natural-ingredient laboratory animal diets.

**Importance:** Published reports of *Clostridium perfringens* isolation from laboratory animal feeds could not be found in the literature. Therefore, we performed a toxin profile screening of our isolated strains of *C. perfringens* to determine which toxinotypes were present in our laboratory animal diets. As studies progress with immunocompromised strains, gnotobiotic models, and animals with perturbed gut flora, the presence of *C. perfringens* could potentially lead to infection, disease and mortality which would substantiate the need to properly eliminate the bacterium and its spores from diets given to high risk animal populations.

## INTRODUCTION

*Clostridium perfringens* is a well-known and widely dispersed gram-positive, non-motile, anaerobic bacterium that ubiquitously inhabits most terrestrial and aquatic environments. Like some other members of the Firmicute phylum, its survivability is greatly enhanced by the bacterium’s ability to produce endospores during periods of environmental stress (1, 2, 3, 4). *C. perfringens* commonly resides in the gastrointestinal (GI) tract of numerous animal species, and opportunistic disease has been widely reported in a variety of domesticated animal species (**Table 1**).

**Table 1:** Pathogenicity of Toxinotypes.

| Disease States in Various Mammals |  |  | Major Toxins |  |  |  | Minor Toxins |  |  |  |
| --- | --- | --- | --- | --- | --- | --- | --- | --- | --- | --- |
|  |  |  | Alpha<br>(CPA) | Beta<br>(CPB) | Epsilon<br>(ETX) | Iota<br>(ITX) | Enterotoxin<br>(CPE) | Beta2<br>(CPB2) | Theta<br>(PFO) | NetB<br>(NetB) |
| Toxinotype | A | • Humans: diarrhea, gas gangrene<br>• Sheep: yellow lamb disease, necrotic enteritis<br>• Poultry: necrotic enteritis<br>• Piglets: enterocolitis, malignant edema | + | — | — | — | — | +/- | +/- | — |
|  | B | • Humans: no reported cases of disease<br>• Sheep/lamb: dysentery, necrohemorrhagic enteritis<br>• Cattle/horses/pigs: enterotoxemia, necrotic enteritis<br>• Goats: enterotoxemia, necrotic enteritis | + | + | + | — | — | +/- | +/- | — |
|  | C | • Humans: food poisoning, necrotic enteritis (pigbel)<br>• Sheep: necrotic enteritis, hemorrhagic dysentery<br>• Calves/kids/foals/piglets: neonatal necrotic enteritis<br>• Poultry: necrotic enteritis | + | + | — | — | +/- | +/- | +/- | — |
|  | D | • Humans: no reported cases of disease<br>• Cattle/sheep/swine/horses: enterotoxemia<br>• Goats: enterotoxemia, colitis<br>• Sheep: enterotoxemia, dysentery, kidney disease | + | — | + | — | +/- | +/- | +/- | — |
|  | E | • Humans: no reported cases of disease<br>• Cattle/sheep/swine/horses: enterotoxemia<br>• Rabbits: enterotoxemia<br>• Calves: hemorrhagic enteritis | + | — | — | + | +/- | +/- | +/- | — |
|  | F | • Humans: diarrhea, CPE-associated nonfoodborne GI disease, gas gangrene<br>• Horses: enteric disease<br>• Dogs: diarrhea, necrotic enteritis | + | — | — | — | + | +/- | +/- | — |
|  | G | • Humans: no reported cases of disease<br>• Avian: gut lesions, necrotic enteritis | + | — | — | — | — | +/- | +/- | + |

(+) toxin is present in all strains of this toxinotype; (-) toxin is absent in all strains of this toxinotype; (+/-) toxin can be either present or absent in strains of this toxinotype References: 2, 3, 14, 15, 18, 19, 31, 32, 33, 36, 40, 79, 80, 81

Not all strains of *C. perfringens* cause disease in animals or humans, and the presence of *Clostridium perfringens* in the intestinal tract usually does not lead to illness. *Clostridium perfringens* typically does not exhibit adherence and invasive properties towards healthy intestinal mucosa, and the development of clinical disease appears to be the result of a complex interaction between host immune status, strain virulence, and other non-specific factors (5). Host stresses leading to abnormal gut microbiota appears to be an important predisposing factor to disease development. Gut microbiota disturbances and vulnerability are known to occur from host antibiotic use, alterations to feeding regimen, overeating and changes to diet (6, 7, 8). Dietary changes known to contribute to clostridial diseases include animal feed with a higher than normal concentration of animal fat, animal protein, fish meal, wheat, barley and indigestible, water-soluble, non-starch polysaccharides (9, 10, 11). *Clostridium perfringens* is the third leading cause of human food poisoning in the world, and disease results from the bacterium’s ability to produce at least 20 known toxins (12). Strains of *C. perfringens* can produce up to four major toxins (alpha, beta, epsilon & iota) as well as numerous other toxins referred to as minor toxins in various combinations. Whereas the presence/absence of all major toxins in a strain of *C. perfringens* determines toxinotype, only a couple of minor toxins, enterotoxin and netB, are correlated with classifying a strain as a particular toxinotype (**Table 1**). Though other toxins produced by *C. perfringens* are referred to as “minor toxins”, clostridial toxins such as enterotoxin, perfringolysin O, netB and beta2 are recognized to be necrotizing and lethal to many animal species (13, 14). The Clo*stridium perfringens* typing system has been recently updated from five toxinotypes (A though E) to seven toxinotypes (A through G). Toxinotype F is a reclassification of enterotoxin-positive (CPE-positive) toxinotype A strains, and toxinotype G strains are produce the necrotic enteritis B-like toxin, netB (15). The following is a brief summary of the major and minor clostridial toxins screened in our study.

### Alpha Toxin (CPA)

Alpha toxin (CPA) is a zinc-dependent phospholipase C enzyme whose activity degrades phosphatidylcholine and sphingomyelin, both components of the eukaryotic cell membrane (**Table 2**). Degradation of the cellular phospholipid membrane results in hemolytic, necrotic and lethal activity (16, 17). *C. perfringens* toxinotype A is the most prevalent toxinotype and is usually present in the normal intestinal flora of humans and other animal species (5). Toxinotype A strains produce only the major toxin α-toxin, and they produce this toxin in higher concentrations than other toxinotypes that possess the CPA gene (18). Alpha toxin is the chief virulent mediator of most diseases caused by *Clostridium perfringens* and is essential in the production of gas gangrene (clostridial myonecrosis) of humans and other mammals including sheep, cattle, goats and horses (3, 5). Experiments have shown alpha toxin to be lethal to mice with a LD_50_ of 3 μg/kg (18, 19).

**Table 2:** Mechanism of Action and Biological Effects of Toxin-Encoding Genes.

| **Toxin** | **Gene** | **Mode of Action** | **Biological Activity** | **Gene Location** |
| --- | --- | --- | --- | --- |
| Alpha ($\alpha$) | cpa | Phospholipase C (plc) and sphingomyelinase | Cytotoxic, hemolytic necrosis, smooth muscle contraction, enteritis | Chromosome |
| Beta ($\beta$) | cpb | Pore-forming activity | Cytotoxic, dermonecrosis, edema, hemorrhagic dysentery, enteritis | Plasmid |
| Epsilon ($\epsilon$) | etx | Pore-forming activity and alteration of cellular membrane permeability | Edema of various organs, smooth muscle contraction, dermonecrosis, blood pressure elevation, enterotoxic | Plasmid |
| Iota ($\iota$) | iap & iep | Ia-mediated ADP-ribosylation of G-actin and pore-forming activity | Disruption of actin cytoskeleton and cell membrane, inhibition of smooth muscle contraction, necrosis | Plasmid |
| Enterotoxin | cpe | Pore-forming activity | Enterotoxic, cytotoxic, erythema, leakage of water and ions, oncosis | Plasmid or Chromosome |
| Beta-2 ($\beta_2$) | cpb2 | Pore-forming activity | Cytotoxic, necrotic and hemorrhagic enteritis, edema, dermonecrosis | Plasmid |
| Theta ($\theta$) | pfo | Perfringolysin and pore-forming activity | Cholesterol-specific hemolysin, tissue destruction, anti-inflammatory | Chromosome |
| NetB | netB | Pore-forming activity | Necrotic enteritis of poultry, hemolytic gut lesions | Plasmid |

*References: 12, 14, 15, 19, 30, 33*

**Table 3:**
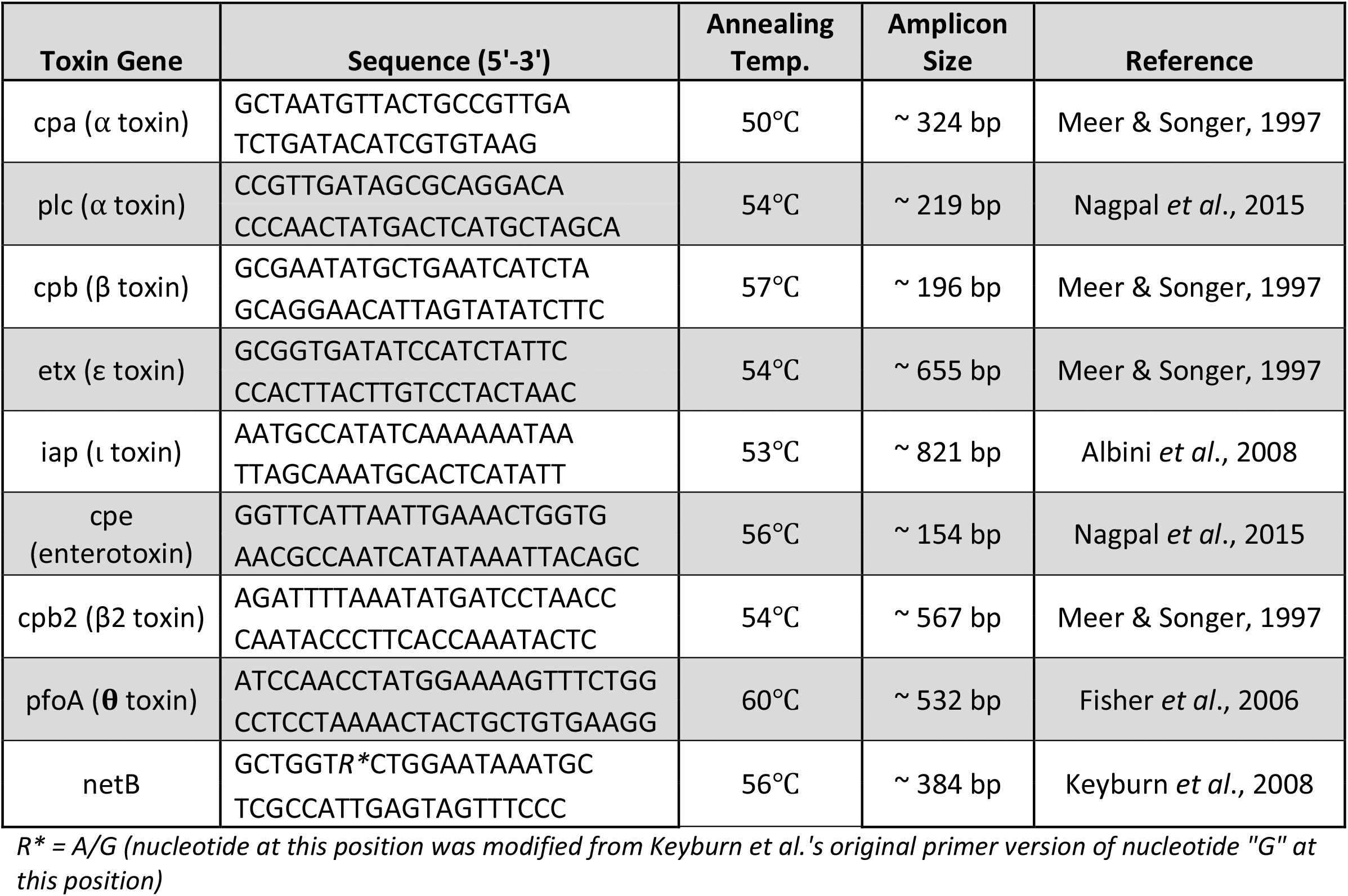
Toxin-Specific Gene Primers.

### Beta Toxin (CPB)

Beta toxin is a pore-forming toxin that disrupts the cellular membrane of susceptible cells. Disease begins in the host’s intestines where absorption of the toxin into the circulatory system often leads to acute neurological signs or sudden death (20, 21, 22). Beta toxin is associated with enteritis necroticans (pigbel) in humans and is responsible for fatal hemorrhagic dysentery in sheep (14, 22) CPB is involved in necrotic enteritis or enterotoxaemia in several domesticated animal species including cattle, goats and sheep (14). Several studies have shown purified beta toxin to be highly lethal to mice with a LD_50_ of 400 ng/kg when injected intravenously (19, 22).

### Epsilon Toxin (ETX)

Epsilon toxin recognizes specific membrane receptors, binds and creates pores within the cellular membrane of susceptible cells creating a rapid and dramatic increase in cellular permeability (23) ETX is produced in the intestine, but the toxin mainly targets distant organs such as the lungs, heart and nervous system (18). Epsilon toxin is considered the major virulence factor of *C. perfringens* toxinotypes B and D and causes blood pressure elevation, increased contractility of smooth muscle, increased vascular permeability and edema of the lungs and brain in many animal species (22, 24, 25). Toxinotype B is very rare in domesticated animals with most cases being reported from the United Kingdom and the Middle East, whereas toxinotype D is the most common cause of enterotoxemia in sheep and goats (22, 26). Behind botulinum toxin and tetanus toxin, epsilon toxin is the third most potent clostridial toxin with a LD_50_ of 100 ng/kg for mice (19, 27).

### Iota Toxin (ITX)

Iota toxin is a binary gene toxin that encodes for two proteins, Ia (enzymatic component) and Ib (binding component). Both proteins are required for pore formation within the cellular membrane of a host cell and are proteolytically activated by trypsin or chymotrypsin (23). The specific role of ITX in disease currently remains unclear (28). Iota toxin causes inhibition of smooth muscle contraction and impairment of endocytosis, exocytosis and cytokinesis (**Table 2**). Only toxinotype E strains produce iota toxin and reported infections associated with iota-toxin are rare, but have been documented in cattle, horses, sheep, swine and rabbits (29, 30, 31, 32, 33). The LD_50_ of iota toxin injected in mice intravenously was determined to be 40 μg/kg (19).

### Enterotoxin (CPE)

Enterotoxin is produced during sporulation of CPE-positive strains and accumulates intracellularly until lysis of the sporangium releases the toxin into the intestinal lumen (5, 36). CPE binds to claudin receptors to form a small complex (**Table 2)**. Oligomerization of several of these complexes on a cell’s surface forms a prepore which eventually inserts itself into the membrane bilayer to create a cation selective pore (37, 38). The formation of cation-permeating pores allows the influx of calcium which leads to cellular death (39). The *cpe* gene is typically located on the bacterium’s chromosome for toxinotype F strains, whereas the *cpe* gene is usually plasmid-based for toxinotype C, D and E strains (36). Studies have shown enterotoxin production is important in toxinotype F strains in causing disease symptoms of food poisoning, abdominal cramps, antibiotic-associated diarrhea and nonfoodborne gastrointestinal diseases in humans and animals (12, 15, 40, 41). The LD_50_ of enterotoxin in mice was estimated to be 81 μg/kg when injected intravenously (19).

### Beta2 Toxin (CPB2)

Beta2 toxin is a more recently discovered pore-forming toxin whose biological activity phenocopies that of CPB, though these two proteins share only 15% amino acid similarity (42). CPB2 toxin is a necrotizing and lethal toxin that can be present in all toxinotypes, and whose gross pathology is characterized by hemorrhage and necrosis of the small and large intestines (18). Beta2 toxin is associated with enteric diseases in domesticated animals including swine, horses, cattle, sheep and goats, as well as enteric diseases in wild animals such as deer, black bears and polar bears (44, 45, 46, 47, 48, 49, 50, 51) The LD_50_ of beta2 toxin for mice was demonstrated to be 0.3 μg/kg by intravenous injection (42).

### Theta Toxin (PFO)

Perfringolysin O (PFO) is a pore-forming toxin that targets the lipid bilayer of eukaryotic membranes and is a member of the cholesterol-dependent cytolysin family of pore-forming toxins (52, 53). Besides *C. perfringens* strains that carry the *cpe* gene on their chromosome, theta toxin is thought to be produced by all other strains of *C. perfringens* (36) Studies have shown that theta toxin acts synergistically with alpha toxin to produce the pathological effects associated with gangrenous lesions (clostridial myonecrosis) in humans and animals (54, 55). Theta toxin also appears to have a synergistic effect with epsilon toxin which may enhance the lethality during enterotoxemia in hosts such as sheep and goats (56). The LD_50_ of theta toxin injected in mice intravenously is 15 μg/kg (19).

### NetB Toxin (NetB)

NetB (<u>N</u>ecrotic <u>E</u>nteritis <u>T</u>oxin <u>B</u>-like) is a β-poreforming toxin that plays an important role in the pathogenesis of avian necrotic enteritis (NE)(57). NetB interacts directly with cholesterol to disrupt the fluidity of the lipid bilayer, allowing an influx of ions such as Na^+^, Cl^-^ and Ca^2+^ which causes cellular death and mucosal necrosis of the small intestine (15, 40). Historically, it was believed alpha toxin was a necessary virulence factor for NE in avian species such as chickens and turkeys, but recent studies have demonstrated that the NetB toxin alone has the ability to cause NE (7, 8, 57, 58). *C. perfringens* strains capable of NE were previously classified as either toxinotype A or C (NE associated with beta toxin), but *C. perfringens* toxinotype A strains producing NetB are now classified as toxinotype G stains (**Table 2**).

### Toxinotyping of *C. perfringens* by Polymerase-Chain Reaction (PCR)

Enzyme-linked immunosorbent assays (ELISAs) have been utilized to toxinotype *C. perfringens* strains, but no ELISA has been developed that can reliably detect the Beta2 (CPB2) toxin and has limited success identifying enterotoxin (CPE) (1, 2). Furthermore, biochemical tests alone cannot distinguish between different *C. perfringens* toxinotypes (26). Recent *C. perfringens* toxinotyping efforts have utilized traditional polymerase-chain reactions or quantitative polymerase-chain reactions (PCR or qPCR) to successfully determine the presence of toxin-associated genes.

Currently, there are no published reports of any *Clostridium* spp. being isolated from natural-ingredient, laboratory animal feeds and their toxinotypes. Here we report the isolation and toxigenic profile of 29 *Clostridium* spp. including *C. perfringens* and five other clostridial species (*C. baratii, C. beijerinckii, C. bifermentans, C. butyricum* and *C. sordellii*) from 10 different laboratory animal diet, including both open and closed formulations, from 4 different commercial feed manufacturers. We also demonstrate that *C. perfringens* can be detected via PCR directly from feed DNA samples. These results demonstrate that opportunistic, pathogenic bacteria, such as *C. perfringens*, are present in unsterilized, natural-ingredient, laboratory animal diets and provide a rationale for feed sterilization prior to use to avoid the introduction of unwanted, potentially pathogenic organisms that may cause disease or introduce unwanted physiological effects.

## RESULTS

### Sequence Identification of Clostridial isolates

From the 23 separate lots of laboratory animal feed tested, we isolated and identified seven different species of Clostridium (*C. perfringens, C. bifermentans, C. butyricum, C. baratii, C. beijerinckii, C. sordellii, C. tertium*) (**Table 4)**.

**Table 4:** *Clostridium* sp. Strains Isolated from Natural Ingredient Laboratory Animal Feed.

| Sample ID | Sample | Feed Manufacturer<sup>a</sup> | Lot<sup>b</sup> | Institute<sup>a</sup> | Species |
| --- | --- | --- | --- | --- | --- |
| ATCC® 12917™ | Positive Control Strain - Toxinotype D | NA | NA | ATCC | *perfringens* |
| ATCC® 3626™ | Positive Control Strain - Toxinotype B | NA | NA | ATCC | *perfringens* |
| ATCC® 8009™ | Positive Control Strain - Toxinotype E | NA | NA | ATCC | *perfringens* |
| ATCC® 27324™ | Positive Control Strain - Toxinotype E | NA | NA | ATCC | *perfringens* |
| Uzal Lab Isolate | Positive Control Strain - Toxinotype G | NA | NA | UC Davis | *perfringens* |
| QA1011-17_F | Diet 1 (NIH-31) | 1 | 1 | NIEHS | *perfringens* |
| QA3081-14_B | Diet 1 (NIH-31) | 1 | 2 | NIEHS | *perfringens* |
| QA1881-14_A | Diet 1 (NIH-31) | 1 | 3 | NIEHS | *perfringens* |
| QA4249-17 | Diet 2 (Closed Formula) | 2 | 1 | NIEHS | *perfringens* |
| QA3578-18_A | Diet 3 (Closed Formula) | 1 | 1 | NIEHS | *perfringens* |
| QA1535-17_A | Diet 3 (Closed Formula) | 1 | 2 | NIEHS | *perfringens* |
| QA411-18_C | Diet 4 (NIH-07) | 1 | 1 | NIEHS | *perfringens* |
| QA1027-18_C | Diet 4 (NIH-07) | 3 | 2 | 2 | *perfringens* |
| QA1025-18_C | Diet 5 (Closed Formula) | 2 | 1 | 2 | *perfringens* |
| QA1026-18_C | Diet 6 (Closed Formula) | 2 | NA | 2 | *perfringens* |
| QA1028-18_C | Diet 7 (NIH-2004 - Swine) | 3 | NA | 2 | *perfringens* |
| QA407-18_A | Diet 8 (Closed Formula) | 2 | NA | 3 | *perfringens* |
| QA408-18_B | Diet 9 (Closed Formula) | 2 | NA | 3 | *perfringens* |
| QA409-18_C1 | Diet 10 (Closed Formula) | 2 | 1 | 3 | *perfringens* |
| QA1787-16_A | Diet 11 (Purified - High-fat) | 4 | NA | NIEHS | *perfringens* |
| QA1011-17_B | Diet 1 (NIH-31) | 1 | 1 | NIEHS | *bifermentans* |
| QA1011-17_C | Diet 1 (NIH-31) | 1 | 1 | NIEHS | *butyricum* |
| QA3081-18_D | Diet 1 (NIH-31) | 1 | 2 | NIEHS | *butyricum* |
| QA4071-14_A | Diet 1 (NIH-31) | 1 | 4 | NIEHS | *bifermentans* |
| QA1024-18_E | Diet 1 (NIH-31) | 1 | 5 | 2 | *baratii* |
| QA3913-17_B | Diet 2 (Closed Formula) | 2 | 2 | NIEHS | *beijerinckii* |
| QA409-18_A | Diet 10 (Closed Formula) | 2 | 1 | 3 | *sordellii* |
| QA1025-18_D | Diet 5 (Closed Formula) | 2 | 1 | 2 | *bifermentans* |
| QA218-16_E | Environmental swab | NA | NA | NIEHS | *tertium* |

<sup>a</sup> Numbers were assigned to commercial feed manufacturers and institutes in place of their name.
<sup>b</sup> Lot numbers assigned solely to indicate separate lots of a given diet.

### Toxin Gene Profiling of *Clostridium perfringens* isolates

As expected, PCR, Sanger sequencing and gel documentation verified the presence of toxin genes in all *C. perfringens* isolates, but not in any other species of *Clostridium* isolated from laboratory animal feed (**Figure 1**). All *C. perfringens* strains isolated from laboratory animal diets possessed the major toxin alpha (CPA), and two of the isolates possessed the major toxin beta (CPB). Nine of the fifteen *C. perfringens* strains isolated from laboratory animal feed (60%) possessed the minor toxin theta (PFO), and one strain possessed the minor toxin beta2 (cpb2). No tested feed possessed the netB gene, with only the positive control *C. perfringens* strain, provided to us from the Francisco A. Uzal Lab at the UC Davis School of Veterinary Medicine, contained the netB gene (**Figure 2**). No other major toxins or minor toxins, such as enterotoxin, was observed in any of laboratory animal feed strains.

**Figure 1:**
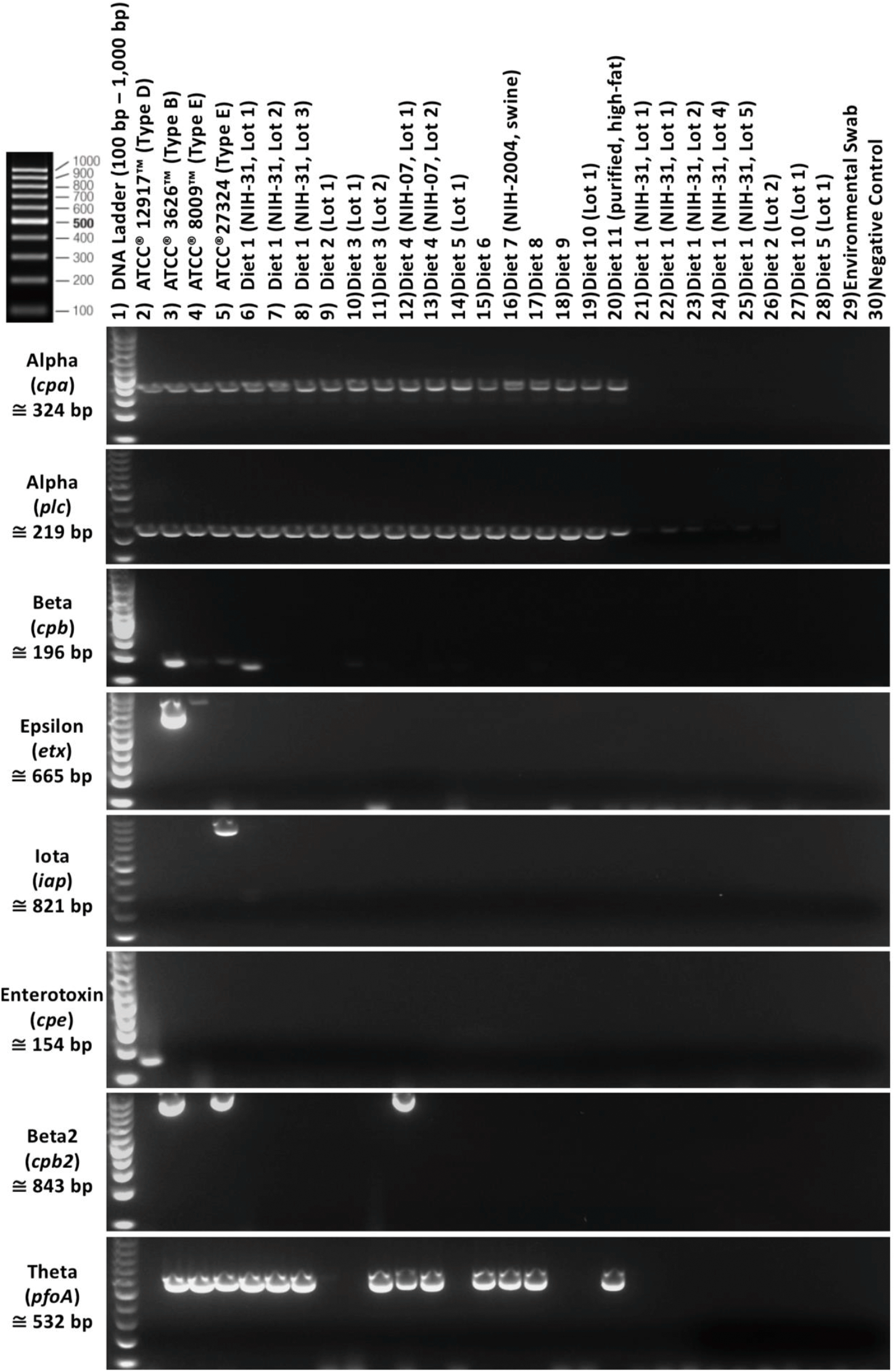
Toxin Gene Profile of *Clostridium sp*. Isolates.

**Figure 2:**
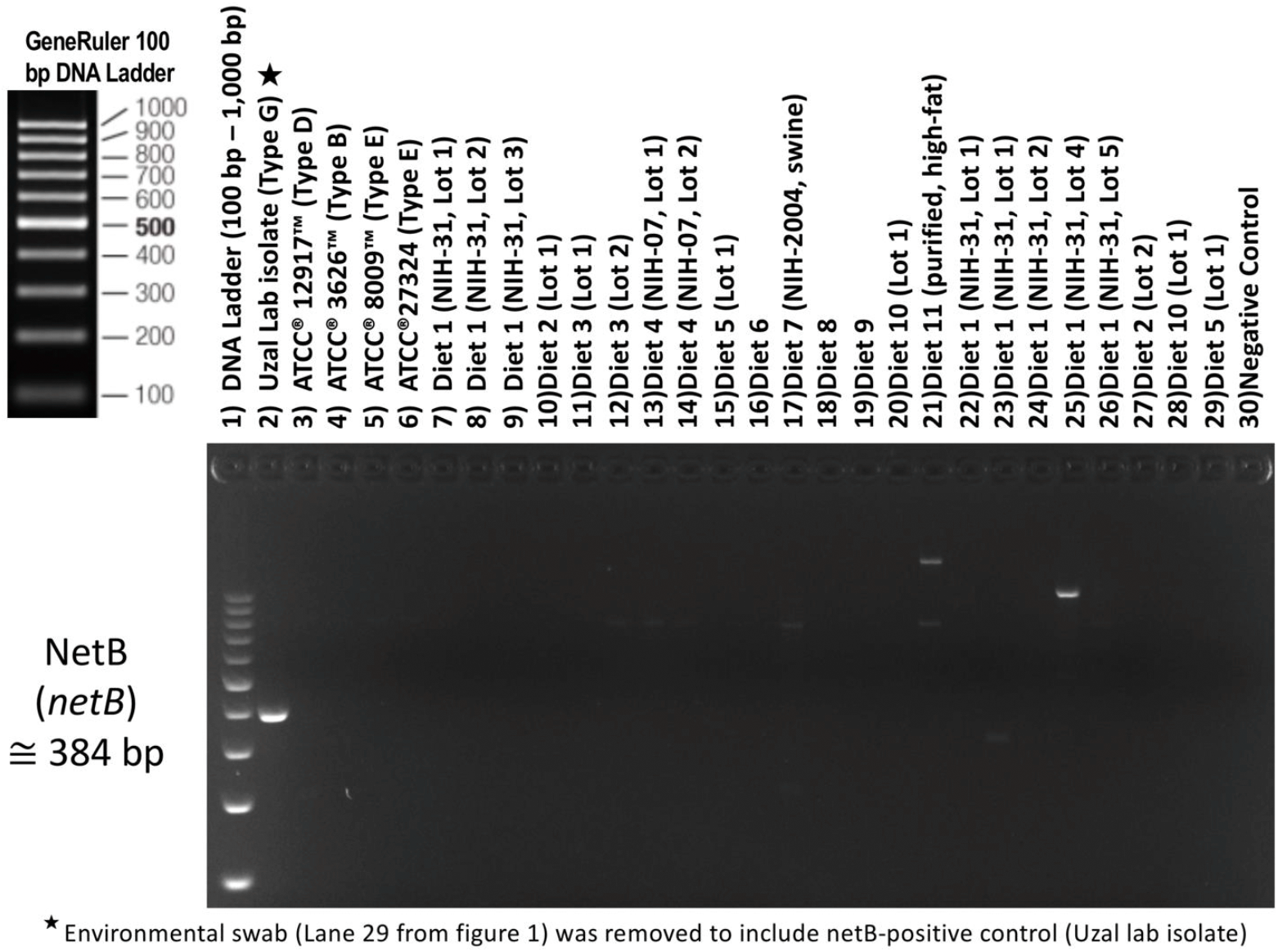
Toxin Gene Profile of *Clostridium sp*. Isolates.

**Figure 3:**
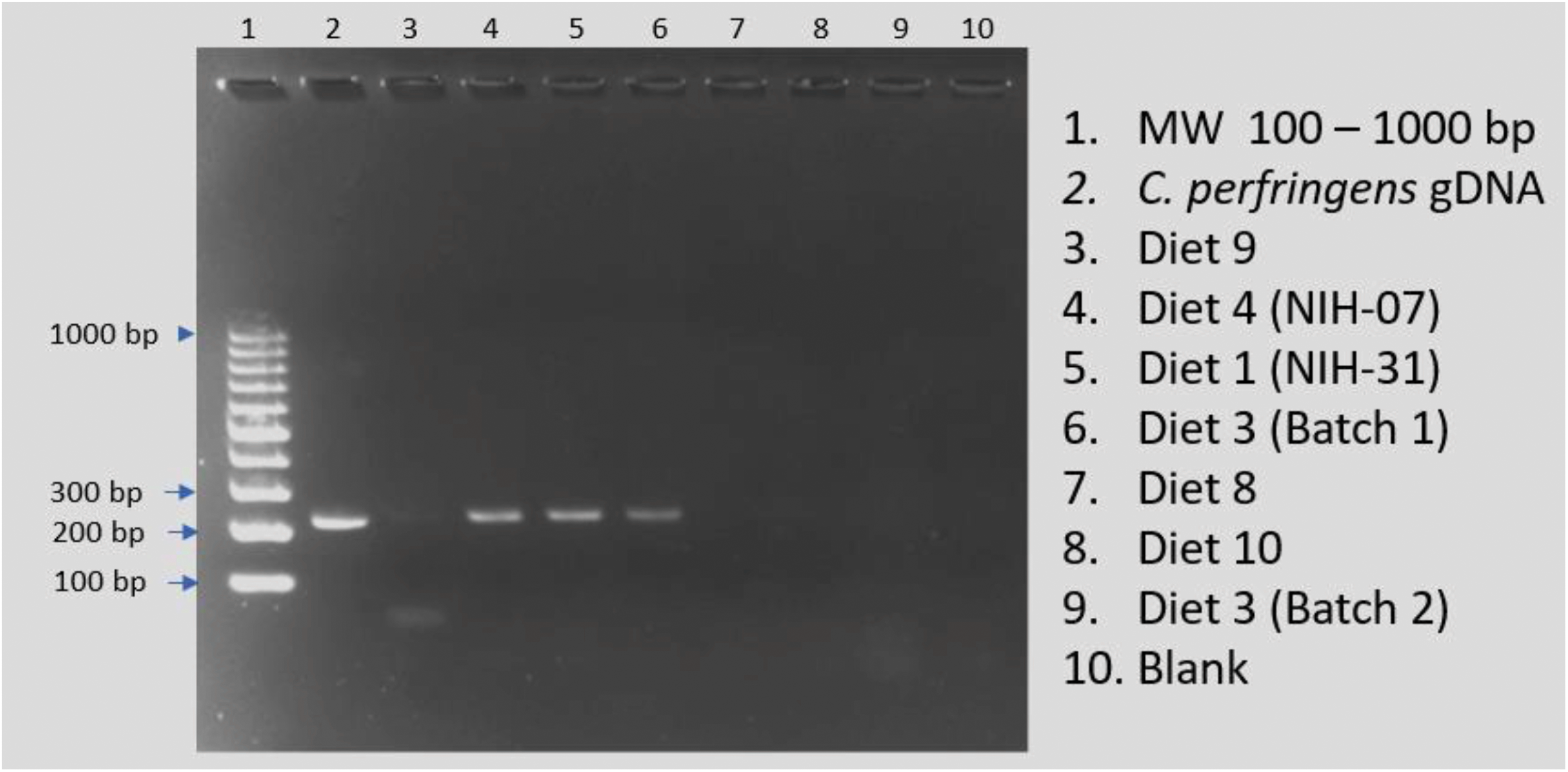
Direct Feed Analysis for *C. perfringens* (α toxin)

However, there were contradictions with three of our four positive control *Clostridium perfringens* strains we used for PCR validation; the toxin profile documentation from the vendor did not correlate with the toxin profile observed from our PCR analysis (**Table 5**). For all three discrepancies (toxin gene absent in positive control strain), another positive control strain containing the toxin of interest was amplified and confirmed that our referenced gene primers worked effectively. Since the toxin gene missing from each of these positive controls (epsilon, iota and enterotoxin) are typically plasmid-based genes, we hypothesize that the plasmid responsible for encoding the respective toxin gene was lost due to continual growth on defined media in a laboratory setting outside an gastrointestinal (GI) environment.

**Table 5:**
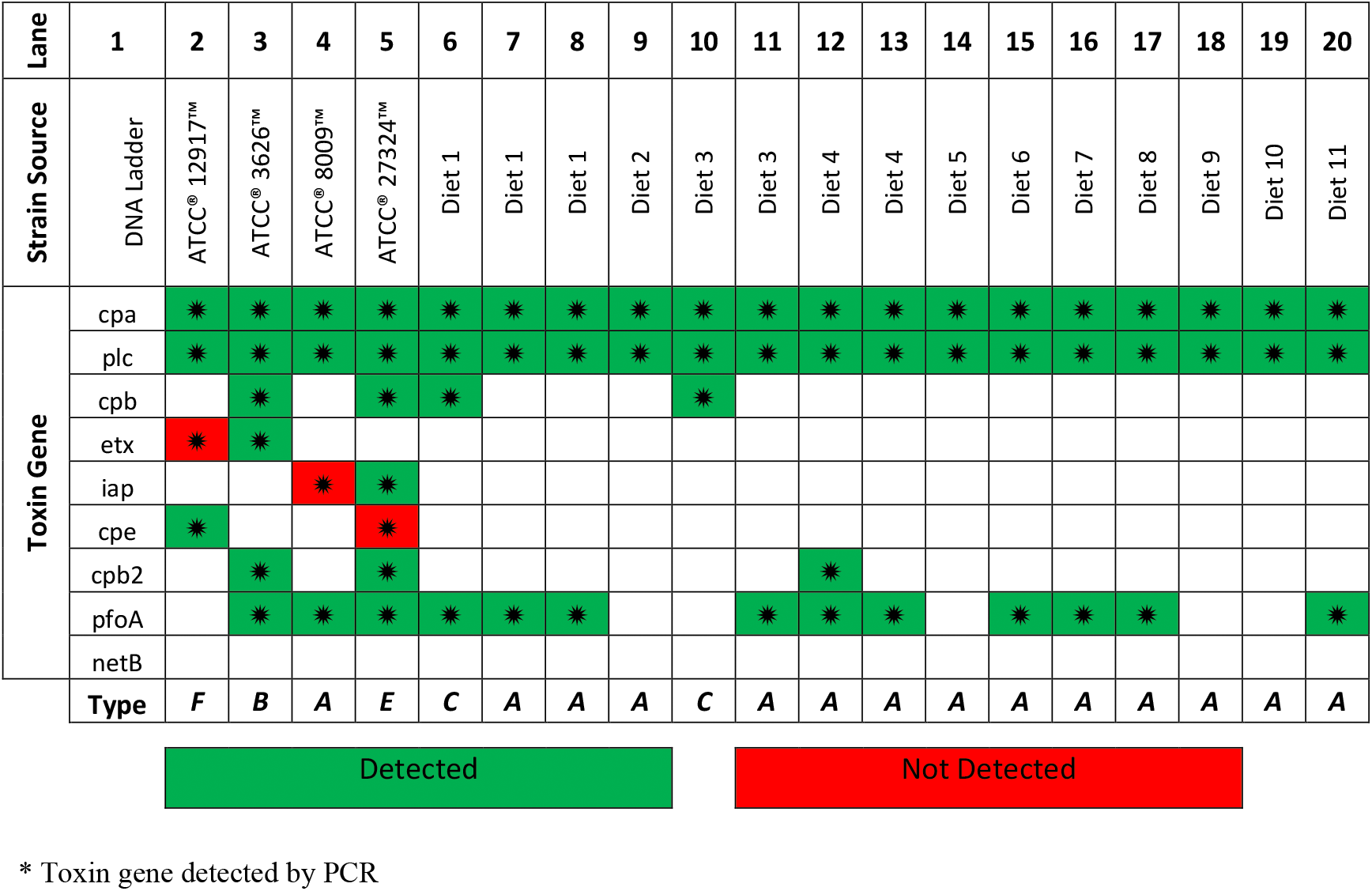
Toxin Profile of *C. perfringens* Strains Isolated from Natural Diets.

### Identification of *C. perfringens* directly from feed

We attempted to identify the presence of *C. perfringens* in animal feed by direct PCR screening for the *plc* (alpha toxin) gene in total gDNA isolated from a subset of tested feeds. All feeds tested for direct PCR were confirmed to possess *C. perfringens* via culture and isolation. Our initial attempts to amplify the gene target directly from feed using the same PCR parameters used on the purified bacterial isolates were unsuccessful (**Table 3**). Using a “touchdown” PCR technique, in which a higher initial annealing temperature (65°C) is used to minimize non-specific amplifications, we were able to correctly amplify the expected 219 bp amplicon from four (4) of the seven (7) diets tested.

## DISCUSSION

Our results demonstrated that almost all of our *Clostridium perfringens* strains were toxinotype A and encoded only for the major toxin alpha which is the major pathogenicity factor in gas gangrene (clostridial myonecrosis) and the chief mediator of many other *C. perfringens*-associated diseases (3). Strains only harboring the major toxin alpha (toxinotype A) are known to produce higher amounts of the alpha toxin than other toxinotypes that also possess a CPA gene. Our results suggest two of the laboratory animal feed strains possess the beta toxin gene whose toxin is known to be fatal in mice at a very low concentration. Though a couple of the beta toxin amplicons observed on electrophoresis gels were faint (low PCR amplification), repeated gel screening and Sanger sequencing of amplicons confirmed their presence to be authentic. We hypothesize that the light banding pattern is the result of only a very small fraction of the bacteria possess a plasmid containing the beta toxin gene. A majority of the *C. perfringens* strains isolated from laboratory animal feed also possessed the theta toxin, and one isolate possessed the beta2 toxin. Both of these minor toxins have been documented to be lethal in certain cases, and studies have shown synergistic interactions and pathogenesis with major toxin alpha (13, 42, 60).

Due to the relatedness of toxin genes in distant relatives of the Clostridium species, such as *C. perfringens, C. sordellii and C. novyi*, the chief mechanisms of such a diverse assortment of toxins appears to be the result of horizontal gene transfer and subsequent independent evolution from strain to strain (61). Amid the broad spectrum of *Clostridial* species, less than 10% produce potent toxins directed against eukaryotic cells. Horizontal gene transfer between *Clostridium* species, as well as non-clostridial species, is well supported by the wide distribution of related pore-forming toxins of the aerolysin family throughout the prokaryotic Kingdom (61). Conceivably these toxin genes evolved from ancestral hydrolytic enzymes by procurement of novel properties such as lipid membrane translocation, pore formation and/or recognition of crucial eukaryotic targets (61).

The ability of *Clostridium perfringens* to transfer toxin-encoding plasmids to neighboring strains of *C. perfringens* is an important element of its pathological progression (19). Only a few of the toxin genes are solely chromosome-encoded (e.g., cpa, pfoA). Most of the toxin genes are solely plasmid-encoded (e.g., cpb, epsilon, cpb2, itx & netB), or, in the case of enterotoxin (cpe), can be either chromosome- or plasmid-encoded (62). Strains of *C. perfringens* can carry up to three different toxin plasmids with each plasmid encoding up to three different toxins (19). Plasmid mobilization (acquisition, loss or exchange between strains) is one of the mechanisms involved in the genetic plasticity of *C. perfringens*, and toxinotype A strains represent the basic toxinotype for which accumulation of toxin-encoding plasmids yield other distinctive toxinotypes (33, 63, 64, 65, 66, 67). Along with antibiotic resistant plasmids, the toxin-encoding plasmids possess a common unique replication region called tcp (transfer of clostridial plasmids) locus (19, 25). The tcp locus can transfer a toxin plasmid from an infecting strain of *Clostridium perfringens* to a normal intestinal flora strain which can intensify and prolong an infection (19).

*Clostridium perfringens* is found ubiquitously in the environment including the digestive tract of healthy animals; therefore, it is difficult to analyze the evolution of pathogenic strains. Additionally, host factors including diet, innate immunity and normal flora greatly impact colonization by foreign bacteria (<u>68</u>, 69). Because few environmental bacteria are known to carry these types of toxin gene variants, it appears evident that strain adaptation to acquire such toxins is suited for survival in a gastrointestinal (GI) environment (61). It has been noted that, unlike commensal strains, pathogenic strains of *C. perfringens* isolated from necrotic enteritis or wound infections show rearrangement of chromosomal regions that include hydrolytic enzymes and toxins which seem to confer selective advantages for colonizing GI environments (70,71). Particular toxinotypes are notably associated with certain animal species such as toxintype C in pigs, toxinotype D in sheep and goats, and toxinotype E in bovine. This specificity suggests certain combinations of chromosomal and plasmid genes are preferential for colonizing the digestive tracts of particular mammals (72, 73).

Because of the bacterium’s ubiquitous nature, almost all food sources, whether animal- or plant-based, can be contaminated with *Clostridium perfringens* (74). Microbial screening for *C. perfringens* has limited effectiveness since positive results are common which indicates very little unless high numbers of *C. perfringens* are enumerated (5). Most sterilization processes, including ionizing radiation, do well at destroying vegetative cells, but do not always effectively eliminate their spores (76). Surviving spores in sterilized feed can germinate and multiply rapidly (77, 78). Proper heating is the most reliable method of spore inactivation, but the required temperature and time is dependent on feed properties such as pH, water content and fat content (75). In compliance with the regulations of animal feed quality for the Republic of Serbia, 50 grams of sample feed must not contain any *C. perfringens* (5). Without toxin genotyping of *Clostridium perfringens* strains and precise calculation of the degree of contamination, risk assessment to animal populations is substandard especially for immunocompromised, gnotobiotic, and germ-free animals.

The genetic profile and pathogenic properties of this bacterium are of significant interest to the Quality Assurance Laboratory. While disease due to *C. perfringens* has not been seen in the rodent population at NIEHS, immunocompromised, gnotobiotic and germ-free animals could be more susceptible to colonization with *C. perfringens* and disease due to toxin production. Therefore, the proliferation of *C. perfringens* from laboratory animal feed could pose a serious risk to studies utilizing these types of rodent models. Since the isolation of *C. perfringens* from laboratory animal diets has not been previously reported, we wanted to develop our own capabilities to rapidly screen incoming diets for *C. perfringens* and effectively evaluate their toxigenic profile via PCR-based assays. While we were able to demonstrate the presence of C. perfringens in laboratory animal diets via direct PCR of total gDNA isolated from feed, our results were not consistent. This is likely due to the concentration of *C. perfringens* in the diet and/or PCR inhibitors remaining in the gDNA isolated from the feed. As such, we caution the use of PCR screening of feed samples as the sole method to screen animal feeds for *C. perfringens*. Based on our findings, *C. perfringens* appears to be a common contaminant of laboratory animal feeds. Almost all of the isolates we identified exclusively fall into the toxinotype A category. Given these findings, we strongly recommend that users sterilize their laboratory animal diets prior to use to prevent the exposure to potentially pathogenic strains of *C. perfringens* which can lead to disease or subclinical alterations in physiology that can affect research outcomes.

## MATERIALS & METHODS

### *Clostridium* spp. Cultivation from Animal Feed and Initial Identification

Twenty-three separate lots of laboratory animal feed were tested. These included several lots of our standard, open-formula NIH-31 rodent feed produced under contract by a commercial source. We also tested two lots of the open formula NIH-07 rodent diet from two different manufacturers; the NIH-2004 open-formula swine diet, four different natural-ingredient, closed-formula rodent diets from two different manufacturers; and one purified, high-fat, rodent diet. For enrichment of *Clostridium* spp., approximately 25 g of each feed sample was aseptically placed into 250 ml of Thioglycolate broth and incubated at 35°C for 24 hours. After incubation, each thioglycolate broth bottle was briefly mixed and streaked onto blood agar plates (BAPs) using sterile cotton tipped applicators. BAPs were then incubated at 35°C under anaerobic conditions using a GasPak™ EZ gas generating system from BD Diagnostics (Franklin Lakes, NJ) inside an anaerobic chamber. After 24-48 hours of incubation, each BAP was examined with suspect colonies isolated onto fresh BAPs and incubated again at 35°C aerobically and anaerobically. Isolates indicating growth only under anaerobic conditions were archived, and a sample of each isolate was shipped to Charles Rivers Laboratories (Wilmington, MA) for identification using matrix associated laser desorption/ionization time of flight mass spectrometry (MALDI-TOF).

### DNA Purification and Quantification

For each isolate identified as a *Clostridium* spp. by MALDI-TOF, a loopful of colony biomass was placed inside a 2-ml tube prefilled with approximately 1200 mg of acid washed, 100 μm zirconium beads (Ops Diagnostics, Lebanon, NJ) along with the initial reagents recommend by the manufacturer of the DNeasy^®^ Blood & Tissue Kit (Qiagen, Hilden, Germany) for “cultured cells”. Next, bead tubes were homogenized using a FastPrep-96™ Homogenizer (MP Biomedicals, LLC, Santa Ana, CA) at max speed for 3 mins. After homogenization, tubes were centrifuged at 8,000 x g for 30 secs and incubated at 56°C for 30 mins. After incubation, tubes were centrifuged at 13,000 x g for 1 min, and each sample’s supernatant was then transferred to a new DNA/RNA-free 1.5 ml microcentrifuge tube. The Qiagen DNeasy^®^ Blood & Tissue Kit’s quick-start protocol was then continued from step 3 until purified DNA was eluted into a new DNA/RNA-free 1.5 ml microcentrifuge tube (step 8). Before downstream analysis, purified DNA was quantified fluorometrically using a DS-11 FX Spectrophotometer/Fluorometer (DeNovix Inc., Wilmington, DE) and a DeNovix^®^ dsDNA Broad Range Assay Kit following the manufacturer’s instructions.Total genomic DNA was isolated from a subset of test diets using the DNeasy PowerMax^®^ Soil kit (Qiagen, Hilden, Germany) to assess our ability to identify *C. perfringens* directly from feed via PCR. Five (5.0) grams of each diet were added to the kit’s 50 ml conical tube along with 0.7 mm garnet beads and 15 ml of the kit’s PowerBead and C1 solutions per kit instructions. The tubes were vortexed for 10 minutes then incubated at 37°C overnight on a shaking tray. The remainder of the kit protocol was followed, and the total genomic DNA was eluted from the Qiagen column using 5.0 ml of C6 solution (elution buffer). The DNA was precipitated with 0.3 M sodium acetate and 10 ml of isopropyl alcohol, washed with 1.5 ml of 75% EtOH, and resuspended in 1.0 ml of DNAse/RNAse-free water. Purified DNA was quantified fluorometrically using a DS-11 FX Spectrophotometer/Fluorometer (DeNovix Inc., Wilmington, DE) and a DeNovix^®^ dsDNA Broad Range Assay Kit following the manufacturer’s instructions.

### Sequence Verification of Cultured Isolates

The bacterial identity of each isolate was verified by 16S rRNA gene analysis using the universal bacterial primers 27F (5’-AGAGTTTGATCMTGGCTCAG-3’, M=C/A) and 1492R (5’-GGTTACCTTGTTACGACTT-3’) (59). Fifty (50) μl polymerase chain reaction (PCR) mixtures were carried out in a T100™ Thermal Cycler (Bio-Rad Laboratories, Hercules, CA) using 100 ng of total DNA template along with AmpliTaq^®^ DNA Polymerase reagents (Applied Biosystems, Foster City, CA) according to the kit’s suggested protocol. PCR amplifications were performed as follows: Initial denaturation at 95°C for 3 min, followed by 35 cycles of 95°C for 30 secs, 55°C for 30 secs, and 72°C for 1 min with a final 5 min extension period at 72°C. 16S rRNA gene amplicons were cleansed using a QIAquick^®^ PCR Purification Kit (Qiagen, Hilden, Germany).

Approximately 50 ng of PCR amplicon template was sent to Genewiz^®^ Inc. (Morrisville, NC) in a premix tube (amplicon + primer) for Sanger sequencing. Sanger sequencing results were trimmed and assembled using CLC Main Workbench 8 (Qiagen, Hilden, Germany) using the default settings. Assembled 16S contigs were identified by uploading sequences into the National Center for Biotechnology Information’s (NCBI) Basic Local Alignment Search Tool^®^ (BLAST^®^) using the ‘16S ribosomal RNA sequences (Bacteria and Archaea)’ database and the top BLAST^®^ score result.

### *Clostridium perfringens* - Toxin Gene Profiling

To screen isolates for various toxin-associated genes, 50 μl PCR reactions were carried out using 10 ng of sample DNA template and toxin-specific gene primers (**Table 3**) along with Platinum™ *Taq* Polymerase (Invitrogen/ThermoFisher, Carlsbad, CA) in accordance with the manufacturer’s guidelines. PCR cycle parameters were as follows: Initial denaturation at 95°C for 3 min, followed by 35 cycles each of 95°C for 30 secs, 50°C - 60°C (annealing temperature for specific toxin gene target) for 30 secs, and 72°C for 1 min with a final 5 min extension period at 72°C. Upon completion of PCR, 20 μl of product was run on a 1.2% agarose gel containing a 1X concentration of GelGreen Nucleic Acid Stain (Candler, NC). As a surety, amplicons were sent to Genewiz^®^, Inc. (Morrisville, NC) for Sanger sequencing, and trimmed results were matched against NCBI’s BLAST^®^ using the ‘nucleotide collection (nr/nt)’ database.

### Identification of *C. perfringens* directly from feed

Fifty (50) nanograms of DNA isolated from feed samples were used in a 50 μl total volume PCR reaction using the *plc* (α) toxin primers listed in **Table 3** and Platinum™ *Taq* Polymerase. Touchdown PCR cycle parameters were as follows: 95°C initial denaturation for 3 min; followed by 10 cycles of 95°C for 30 sec, 65°C for 30 sec with a 0.5°C decrease/cycle, and 72°C for 30 sec; followed by 25 cycles of 95°C for 30 sec, 60°C for 30 sec, and 72°C for 30 sec; and ending with a single 72°C extension for 3 min. Twenty (20) μl of each reaction was run on a 2.0% agarose gel containing 1X GelGreen Nucleic Acid Stain.

## ACKNOWLEDGEMENTS

We would like to thank, Mr. Dennis Barnard, NIH – Division of Veterinary Resources, and Mr. Ned Collins, Alpha-Omega, Inc. – US EPA contractor, for assistance, and the NIEHS Summer Internship Program for funding a portion of this project. We would also like to express our gratitude to Francisco A. Uzal and Juliann Beingesser for providing our lab a netB positive control strain to validate our netB PCR assay.

## DATA AVAILABILITY

As is the policy for NIH employees, the peer-reviewed article will be made publicly available on PubMed Central immediately upon acceptance for publication.

